# White-Matter BOLD Encoding Beyond Marginal Connectivity

**DOI:** 10.64898/2026.07.08.737282

**Authors:** Muwei Li, Zhaohua Ding, John C Gore

## Abstract

Functional MRI studies have traditionally focused on gray matter, whereas white-matter BOLD signals have often been treated as weak or artifactual. Recent work suggests that white-matter BOLD fluctuations contain reproducible functional information, but most gray-to-white matter analyses rely on marginal functional connectivity, which cannot separate pairwise coupling from shared variance among distributed cortical systems. Here, we used a multivariate cortical encoding framework to test whether spontaneous white-matter BOLD activity can be predicted from distributed cortical gray-matter activity and whether this predictive structure reveals organization beyond marginal connectivity. Resting-state fMRI data from 81 Human Connectome Project young adult participants were analyzed using a strict white-matter mask with no overlap with cortical predictors. For each white-matter voxel, time series from 400 Schaefer cortical parcels were used to predict held-out white-matter BOLD signals with nested leave-one-run-out ridge regression. Cortical activity modestly but reliably predicted white-matter BOLD dynamics, demonstrating consistent cross-validated prediction accuracy across a broad spatial extent of the white matter. Ridge beta fingerprints strongly recapitulated marginal functional connectivity fingerprints, indicating a shared functional backbone, but their first gradients diverged reproducibly. This beta-FC divergence axis organized FC-adjusted prediction residuals and remained robust after controlling for gray-matter proximity, mask-boundary distance, white-matter prevalence, temporal signal variability, spatial coordinates, and spatial autocorrelation. The high-divergence end showed relatively low marginal FC but high FC-adjusted prediction residuals and was enriched for posterior thalamic/optic-radiation and posterior corona-radiata anatomy. These findings suggest that multivariate cortical encoding reveals a tract-organized dimension of white-matter functional coupling not captured by pairwise connectivity alone.

## INTRODUCTION

Functional magnetic resonance imaging (fMRI) has fundamentally advanced our understanding of the macroscopic functional organization of the human brain. Historically, however, most fMRI research has been gray matter (GM) centric, focusing on blood-oxygen-level-dependent (BOLD) signals in the cerebral cortex and subcortical nuclei (Logothetis et al., 2001). White matter (WM) BOLD signals were long treated as artifactual or as nuisance variation, in part because WM has lower vascular density, reduced cerebral blood flow, and distinct metabolic properties relative to GM (Gawryluk et al., 2014; Gore et al., 2019; Rostrup et al., 2000). However, growing evidence suggests that WM BOLD signals contain meaningful functional and biological information (Ding et al., 2018, 2013; Gawryluk et al., 2011; Li et al., 2019). Spontaneous low-frequency fluctuations in WM are not random; they exhibit distinct spectral properties (Li et al., 2021), show structural-functional coupling, and form reproducible functional patterns related to global neurophysiological activity (Ding et al., 2013; Schilling et al., 2019). These findings have shifted WM fMRI from a purely methodological concern toward an important target for studying whole-brain functional organization.

To date, resting-state investigations of WM functional organization have relied heavily on marginal functional connectivity (FC) between WM and GM. These approaches typically estimate zero-lag Pearson correlations between individual WM voxels or tracts and distributed cortical parcels or large-scale resting-state networks (Gao et al., 2023, 2020; Li et al., 2024a, 2023). This strategy has revealed intrinsic WM functional networks that topologically mirror overlying GM systems, including default mode, frontoparietal, and somatomotor networks (Ji et al., 2019; Peer et al., 2017). Variation in marginal gray-to-white FC profiles has also been linked to cognition, development, and neurological disease (Wang et al., 2022; Xu et al., 2024). Nevertheless, marginal FC is limited by broad cortical synchrony, spatial autocorrelation, and collinearity among large-scale networks. Because distributed cortical regions often fluctuate together, a pairwise WM-GM correlation cannot determine whether an apparent association reflects a specific cortical contribution or shared variance with other synchronized cortical systems (Cole et al., 2013; Smith et al., 2011). Thus, marginal FC provides an important but incomplete description of gray-to-white functional coupling.

Multivariate regression and encoding models provide a principled way to model signals in the presence of pervasive predictor collinearity. In task fMRI, voxelwise encoding models with regularization, including ridge and Lasso regression (Hilt et al., 1977; Tibshirani, 1996), have been used to map complex and highly correlated stimulus features, such as natural vision, semantic content, and language, onto cortical BOLD responses (Huth et al., 2016, 2012; Kay et al., 2008; Naselaris et al., 2011). Ridge regression is particularly useful when predictors are strongly correlated because its L2 penalty stabilizes model estimates and distributes predictive weights across related features. In resting-state fMRI, however, multivariate modeling has more often been used in decoding or prediction frameworks, such as connectome-based predictive modeling, where distributed connectivity patterns are used to predict behavioral traits or cognitive scores (Finn et al., 2015; Rosenberg et al., 2016; Shen et al., 2017). By contrast, using distributed resting-state GM activity as an internal conditional encoding model to predict localized BOLD dynamics remains less explored, especially for mapping WM functional architecture.

A multivariate approach is particularly relevant for WM because recent work suggests that WM engages dynamically in whole-brain functional organization rather than serving only as a passive structural relay (Li et al., 2024b, 2020). These findings imply that WM signals may reflect many-to-one relationships with distributed cortical systems, a setting in which pairwise FC can obscure conditional predictive structure. Inspired by this view and by the success of encoding models in cortical fMRI, we propose that modeling how distributed cortical GM activity predicts local WM signals can complement existing marginal engagement and connectivity metrics. Rather than treating WM-GM interactions as isolated pairwise correlations, a predictive encoding framework can assess cortical contributions after accounting for shared variance among cortical predictors.

In the present study, we investigated whether spontaneous BOLD activity in human WM can be predicted from cortical GM activity, and whether this predictive relationship contains functional organization beyond conventional marginal FC. Using high-quality resting-state fMRI data from 81 Human Connectome Project (HCP) young adult participants, we modeled strict WM voxel time series as a function of 400 distributed cortical parcels (Schaefer et al., 2018) using nested, leave-one-run-out cross-validated ridge regression. We extracted high-dimensional beta fingerprints from this conditional encoding model and directly compared them with traditional marginal FC fingerprints. We expected cortical activity to predict WM BOLD dynamics reliably but modestly. We further hypothesized that ridge beta fingerprints would share a strong functional backbone with marginal FC, while the divergence between the two, conceptualized as a beta-FC divergence axis, would reveal a reproducible conditional encoding dimension. Finally, by controlling for spatial autocorrelation, GM proximity, and local signal quality, we tested whether this divergence axis organizes FC-adjusted prediction residuals, reflects cortical network contributions, and aligns with WM tract anatomy. This framework aims to separate marginal gray-to-white connectivity from conditional cortical encoding, providing a more nuanced view of how distributed cortical activity relates to WM functional dynamics.

## METHODS

### Ethics Statement

The data involved in this research are publicly available and have been previously approved for use by the Washington University Institutional Review Board. All participants provided written informed consent to participate in this study. The authors did not collect any new data involving human participants.

### Dataset

We used resting-state fMRI data from the Human Connectome Project Young Adult (HCP-Y) cohort (Van Essen et al., 2013). The initial sample was the classic HCP unrelated 100-subject set (Barch et al., 2013). Subjects were retained only if all four 3T resting-state fMRI runs (rfMRI_REST1_LR, rfMRI_REST1_RL, rfMRI_REST2_LR, and rfMRI_REST2_RL) had complete physiological recordings (Physio_log.txt), complete motion-regressor files (Movement_Regressors_dt.txt), and no preprocessing errors during covariate generation. Nineteen subjects failed these criteria and were excluded, leaving 81 participants for analysis. The final sample included 44 females and 37 males; 12 participants were aged 22-25 years, 34 were aged 26-30 years, and 35 were aged 31-35 years. Sex was recorded by self-report in the HCP dataset, and no sex- or gender-based analyses were conducted.

HCP-Y imaging protocols are detailed elsewhere (Van Essen et al., 2012). Briefly, data were acquired on 3T Siemens Skyra scanners. Resting-state fMRI was acquired using multiband gradient-echo EPI sequences across two sessions, with each session containing two runs acquired with opposing phase-encoding directions. Each resting-state run lasted 14 minutes and 33 seconds and contained 1200 volumes, with TR = 720 ms, TE = 33.1 ms, and 2 mm isotropic spatial resolution. Cardiac and respiratory physiological signals were recorded concurrently during fMRI acquisition. T1-weighted images were acquired using a single-echo MPRAGE sequence with TR = 2400 ms, TE = 2.14 ms, and 0.7 mm isotropic resolution. T2-weighted images were acquired using a 3D T2-SPACE sequence with TR = 3200 ms, TE = 565 ms, and 0.7 mm isotropic resolution.

### Preprocessing

We started from HCP minimally preprocessed resting-state fMRI data (Glasser et al., 2013). The HCP structural and functional preprocessing included nonlinear registration of T1-weighted images to MNI space using FNIRT (Jenkinson et al., 2012), FreeSurfer-based anatomical processing (Dale et al., 1999), correction of fMRI head motion and susceptibility-related distortions, and registration of functional data to MNI space. We then applied project-specific preprocessing to each retained resting-state run. Physiological covariates were generated from the HCP Siemens physiological log files using TAPAS PhysIO (Kasper et al., 2017), including RETROICOR terms (Glover et al., 2000), respiratory volume per time, and heart-rate variability; HCP motion regressors were appended to the physiological regressors. For each run, voxelwise time series were regressed on the physiological and motion covariates, a linear trend, and an intercept. The residual time series were then band-pass filtered at 0.01-0.1 Hz, and voxel means were added back before writing the preprocessed 4D images. Individual WM and cortical GM masks were generated from HCP FreeSurfer wmparc segmentations. The group WM95 mask was defined as voxels classified as WM in more than 95% of retained subjects, and the group GM mask threshold was selected to ensure zero overlap with the group WM mask. The final analysis masks were in the same image space, contained 32,445 strict WM voxels and 149,268 GM voxels, and had no GM-WM or Schaefer-atlas-to-WM overlap.

### Cortical and WM time-series extraction

For each retained run, cortical GM predictor time series were extracted from the Schaefer-400 atlas (Schaefer et al., 2018) by averaging the preprocessed BOLD signal across all voxels assigned to each cortical parcel. WM target time series were extracted from the strict group WM95 mask. Before extraction, WM data were smoothed within the WM mask using 4 mm full- width-at-half-maximum mask-normalized Gaussian smoothing. Specifically, each volume was multiplied by the binary WM mask, smoothed, and divided by a smoothed mask denominator so that smoothing remained constrained to the analysis mask. This step was used to improve local WM signal-to-noise while avoiding spillover from outside the strict WM mask.

### Voxelwise cortical encoding model

The primary analysis used voxelwise ridge regression to predict each WM voxel time series from the 400 cortical parcel time series. Model performance was evaluated using nested leave-one-run-out cross-validation. In each outer fold, one run was held out for testing, and the remaining three runs were used for training and inner-loop model selection. The ridge penalty was selected only within the outer-training data using inner run-level cross-validation and a deterministic sample of WM voxels. The selected model was then refit to the outer-training runs and evaluated on the held-out run. Ridge models were fit after standardizing predictors and targets within each training fold. In its standard form, ridge regression estimates coefficients by minimizing the penalized least-squares objective:

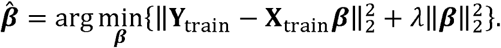

In the present implementation, the ridge penalty was normalized by the number of training time points, yielding the closed-form solution:

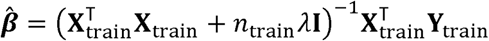

where X_train_ and Y_train_ denote the standardized cortical predictor matrix and WM target matrix, n_train_ is the number of training time points, λ is the ridge penalty, and I is the identity matrix. This sample-size normalization made the ridge penalty comparable across folds. Prediction accuracy was summarized voxelwise as cross-validated R² and prediction correlation across the held-out time points. Subject-level mean R² and the fraction of voxels with positive R² were used as summary measures.

Two model-sensitivity analyses were used to contextualize the primary 400-parcel zero-lag model. First, a reduced 17-network model was fit by averaging Schaefer parcels within their 17-network labels (Thomas Yeo et al., 2011) and repeating the same nested cross-validation procedure. Second, a lagged ridge model was fit using cortical predictors shifted across a fixed set of temporal lags. Its performance was compared with a zero-lag model refit on the identical edge-trimmed time points. These sensitivity analyses were used to determine whether fine-grained cortical predictors and temporal lag structure changed the level or spatial extent of WM prediction.

### Group-level prediction maps and circular-shift null validation

Subject-level R^2^ maps were written back to the strict WM mask and combined across subjects to generate group means, median, t-statistic, FDR-corrected, and positive-consistency maps. Group-level inference on mean R^2^ used voxelwise one-sample tests across subjects, followed by false-discovery-rate correction across WM voxels.

To evaluate whether the observed prediction exceeded a temporal autocorrelation-preserving null, circular-shift null models were generated within each subject. For each null iteration, the WM target time series were circularly shifted within run by at least 20 s, while preserving the original cortical predictors and the fold-specific lambdas selected in the observed model. Null R^2^ maps were accumulated across 1000 iterations per subject, and voxelwise p-values were computed from the fraction of null R^2^ values exceeding the observed R^2^. These p-values were FDR corrected within each subject to summarize the spatial extent of above-null prediction.

### Ridge beta fingerprints, marginal FC fingerprints, and model comparison

For downstream coefficient analyses, the primary cross-validated prediction model and the beta fingerprint analyses used separate lambda policies. Cross-validated R^2^ was always taken from the subject- and fold-specific nested models. To make beta coefficients more comparable across subjects, final beta fingerprints were refit using all four runs and a shared lambda defined as the geometric median of the subject-level final lambdas. These shared-lambda beta estimates were used only for fingerprint comparison and gradient analyses, not for out-of-sample prediction.

For each subject, each WM voxel was represented by a 400-dimensional standardized ridge beta fingerprint. In parallel, a marginal FC fingerprint was computed as the Pearson correlation between each WM voxel time series and each cortical parcel time series after concatenating all runs. Beta and FC fingerprints were compared at four levels: signed 400-parcel profile correlation, absolute-weight profile correlation, 17-network composition correlation, and overlap among the strongest 5 percent of parcel weights. Group-level comparisons were computed from group-average beta and FC profiles, and subject-level distributions were computed within each participant.

Cortical network contribution maps were also computed using leave-one-network-out predictive ablation. For each Yeo 17-network predictor group, model performance was recomputed after removing cortical predictors of that network, and the change in R^2^ relative to the full model was used as the network contribution score. Because cortical networks are correlated, these Delta R^2^ values were interpreted as predictive ablation scores rather than additive or causal contributions.

### Fingerprint stability and beta-FC gradient construction

The reliability of beta and FC fingerprints was assessed using two independent split schemes. The session split compared REST1 runs with REST2 runs, whereas the balanced split compared REST1_LR plus REST2_RL with REST1_RL plus REST2_LR. For each split, beta fingerprints were refit using the shared lambda, and FC fingerprints were recomputed from the corresponding runs. Voxelwise fingerprint stability was defined as the correlation between the two independent 400-dimensional profiles.

Low-dimensional fingerprint organization was estimated using BrainSpace gradient mapping with a normalized-angle kernel and diffusion embedding (Vos de Wael et al., 2020). Because ridge beta fingerprints are signed, sparsity thresholding was set to zero so that negative coefficients were preserved. Split gradients were aligned using Procrustes alignment, and matched gradient components were compared across splits. The first beta and FC gradients were used to define the dominant low-dimensional axes of conditional beta fingerprints and marginal FC fingerprints, respectively.

A fixed beta-FC G1 template was constructed from the independently estimated session-split gradients. The FC G1 template was sign-aligned to the beta G1 template because the gradient sign is arbitrary. Linear projection weights were then learned from group-average beta and FC fingerprints to their respective G1 templates. The beta and FC fingerprints of each subject were projected onto these fixed templates, z-scored, and differenced to produce a subject-level beta-minus-FC G1 map. Group-level inference on this difference map used voxel-wise one-sample tests and sign-flip max-T family-wise-error correction across WM voxels.

### Local prediction residuals and beta-FC divergence

To test whether cortical encoding captured local WM organization beyond marginal FC strength, we computed local maps of prediction accuracy and FC magnitude. Within each subject, voxelwise R^2^ values were z-scored across WM voxels. Marginal FC strength was summarized as the root-mean-square of each voxel’s 400-dimensional FC profile and was also z-scored across WM voxels. A local R^2^ residual map was then obtained by regressing z-scored R^2^ on z-scored FC strength within the subject. This residual quantified where cortical-to-WM prediction was higher or lower than expected from local marginal FC magnitude.

Group local R^2^ residual maps were computed by averaging subject residual maps. Voxelwise inference used subject-level sign-flip testing with max-T FWER correction. In addition, positive and negative residual clusters were formed separately using a two-sided cluster-forming threshold, and cluster significance was assessed with signed cluster-mass sign-flip permutations. The spatial association between local R^2^ residuals and beta-minus-FC G1 differences was summarized using voxelwise density plots and map correlations.

### Spatial, signal-quality, and spatial-autocorrelation controls

Several voxelwise nuisance maps were constructed to test whether the local residual and beta-FC G1 effects could be explained by simple anatomical or signal-quality factors. These covariates included distance to the group GM mask, distance to the WM mask boundary, subject-level WM mask prevalence, mean extracted WM temporal standard deviation, and spatial coordinates with low-order polynomial and interaction terms. Distance maps were computed in millimeters within the common image space.

For each subject, local R^2^ residual maps and beta-minus-FC G1 difference maps were residualized against the nuisance design. A fully controlled prediction map was also computed by residualizing z-scored R^2^ against local FC strength and the same spatial nuisance covariates. Controlled group maps were inferred using sign-flip max-T FWER correction. Joint controlled effects were summarized as voxels significant for both the controlled R^2^ residual and controlled beta-FC G1 difference, with sign agreement used to determine whether the two controlled effects converged spatially.

Because WM maps are spatially autocorrelated, we also used a volumetric spatial-block permutation null. WM voxels were assigned to 12-mm spatial blocks based on template coordinates. In each permutation, the beta-FC G1 map or significance mask was permuted at the block level, preserving local spatial structure within blocks. This null was used to test the observed residual-G1 map correlation and the observed same-sign joint voxel count.

### Anatomical, network, hemispheric, and validation summaries

To summarize the anatomical organization of the beta-FC divergence axis, WM voxels were divided into ten equal-count bins ordered by the beta-minus-FC G1 axis. Within each bin, we summarized mean zR2, local FC strength, FC-adjusted R^2^ residuals, nuisance-controlled residuals, cortical network contribution, distance and prevalence covariates, and tract composition. Tract composition was defined by direct overlap with the JHU WM atlas (Mori et al., 2009; Oishi et al., 2008). JHU tract-level summaries were also computed by averaging local residual and beta-FC G1 values within each tract; a 4-mm proximity analysis was retained as a sensitivity check.

Hemispheric organization was assessed by pairing left-hemisphere WM voxels with their right-hemisphere mirror coordinates in template space. For each major map, left-right mirror correlations, absolute left-right differences, and signed left-right differences were computed. These analyses were used to describe bilateral organization rather than to define primary statistical significance.

Two validation analyses were used to assess reproducibility. First, local R^2^ residuals were estimated separately in REST1 and REST2 using one phase-encoding run to predict the other within each session, and REST1-REST2 map correlations and same-sign conjunctions were computed. Second, beta-FC G1 projections learned from one session were applied to subject fingerprints from the opposite session, and the two cross-session directions were compared. Behavioral prediction analyses using global encoding features and local JHU-tract summaries were treated as exploratory sensitivity analyses. They used nested cross-validation with age, sex, and mean motion as unpenalized confounds, and incremental prediction beyond confounds was tested with Freedman-Lane residual permutations (Freedman and Lane, 1983).

## RESULTS

### Strict WM prediction from cortical activity

We first verified that the anatomical inputs provided a strict gray-white separation for cortical-to-WM prediction. The Schaefer-400 cortical atlas and group WM95 mask occupied the same image space, with 128,801 cortical atlas voxels, 149,268 GM voxels, and 32,445 strict WM target voxels. The atlas-WM and GM-WM overlap counts were both zero, confirming that cortical predictors and WM target voxels were anatomically separated. The final analysis included 324 resting-state runs from 81 participants (Fig. S1).

Using these inputs, cortical parcel activity predicted held-out WM BOLD signals above zero across a broad portion of the WM mask. In the primary zero-lag 400-parcel ridge model, the mean subject-level cross-validated R^2^ was 0.0199 +/- 0.0177, and an average of 70.3% +/- 7.1% of WM voxels had positive held-out R^2^. The group-mean R^2^ map showed spatially structured prediction rather than a uniform low-amplitude effect, indicating that distributed cortical activity explained a modest but reliable component of WM BOLD variance (Fig. 1a-d).

**Figure 1.**
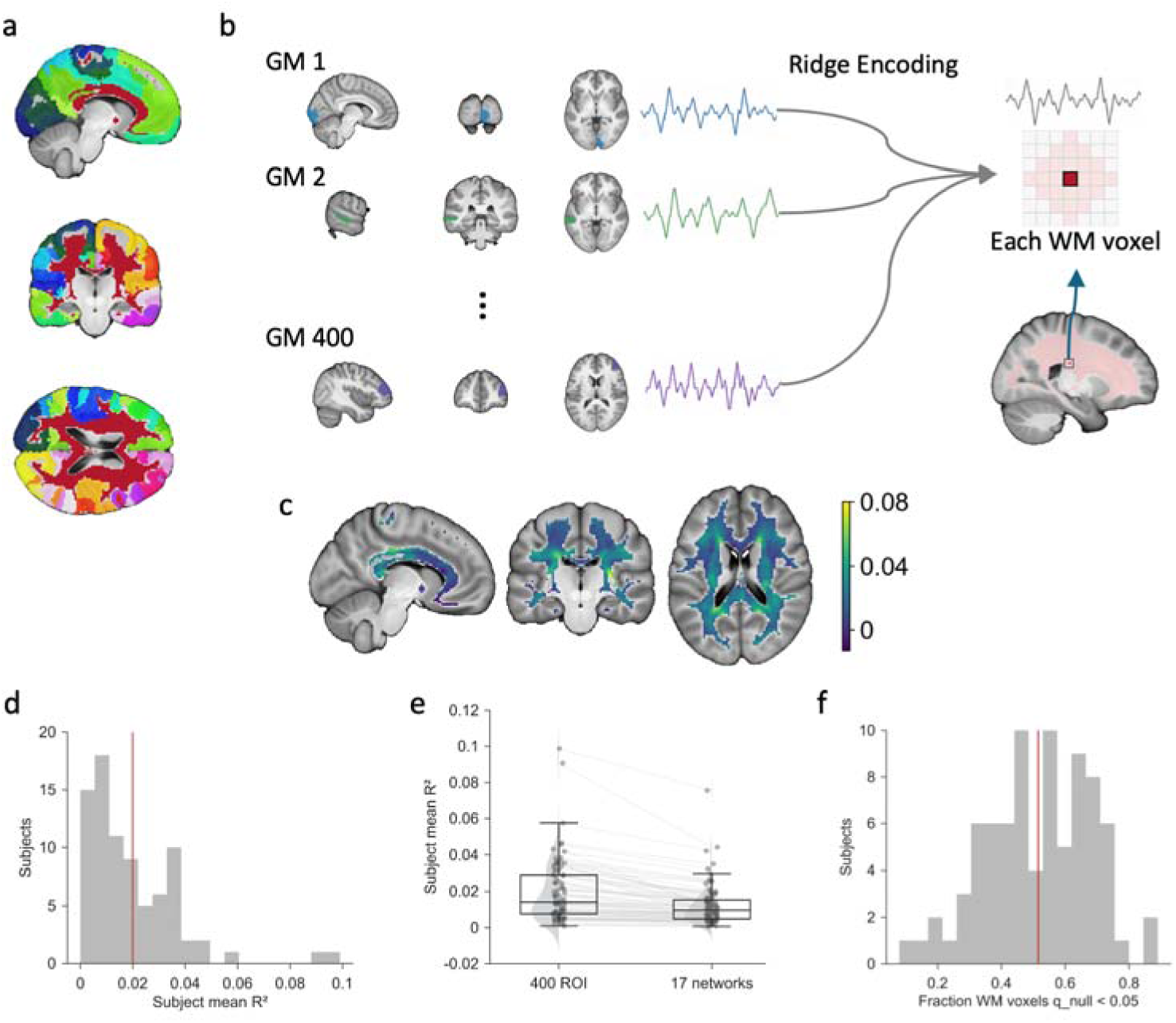
Cross-validated cortical encoding of WM BOLD activity. (a) Anatomical definition of cortical predictors and WM targets. Schaefer-400 cortical GM parcels are shown with distinct atlas colors, and the strict WM95 target mask is shown in red. (b) Ridge-encoding model schematic. For each WM voxel, time series from 400 cortical GM parcels were used as predictors in a ridge-regression model. The regularization parameter was selected using nested cross-validation, and prediction accuracy was evaluated on held-out data. (c) Group-mean voxel-wise cross-validated R² for the zero-lag 400-parcel encoding model, showing the spatial distribution of WM BOLD variance predicted from cortical GM activity. (d) Distribution of subject-level mean WM R² values. The red vertical line indicates the sample mean. (e) Within-subject comparison of prediction accuracy between the Schaefer-400 parcel model and a reduced 17-network model. Points and boxes show subject-level distributions, and grey lines indicate paired values from the same subject. (f) Circular-shift null validation. The histogram shows, for each subject, the fraction of WM voxels with q_null < 0.05; the red vertical line indicates the sample mean.

Model-sensitivity analyses supported the use of fine-grained cortical predictors while showing that the main conclusion did not depend on a single modeling variant. The 400-parcel model outperformed the reduced 17-network model by a mean within-subject difference of 0.0074 R^2^, indicating that parcel-level cortical detail contributed to WM prediction (Fig. 1e; Fig. S2; Fig. S4). The lagged ridge model produced only a small average increase over the matched zero-lag model (mean difference = 0.0020 R^2^), suggesting that the zero-lag model captured the dominant predictive structure used in the main analyses (Fig. S2; Fig. S4). Subject-specific final lambdas were centered near the shared lambda used for downstream beta refits (median = 36.5; shared lambda = 36.5), supporting a common shrinkage reference for coefficient comparisons (Fig. S2).

A circular-shift null model confirmed that above-zero prediction was not explained simply by temporal autocorrelation. For each participant, WM target time series were shifted within run while preserving the original cortical predictors and selected lambdas; observed prediction was then compared with the resulting null distribution. Many subjects retained WM voxels surviving q_null < 0.05, and the spatial consistency map showed recurrent above-null prediction across subjects (Fig. 1f; Fig. S3).

### Ridge beta fingerprints share an FC backbone but define a distinct stable gradient

We next asked whether multivariate ridge beta fingerprints simply reproduced marginal GM-WM functional connectivity (FC) or provided additional organization. Across WM voxels, ridge beta and FC fingerprints were strongly related: the group signed profile correlation was 0.917, the absolute profile correlation was 0.781, the 17-network composition correlation was 0.794, and the top-ROI overlap was 0.749. These relationships were also consistent at the subject level, with a mean signed beta-FC profile similarity of 0.948 +/- 0.018. Thus, ridge coefficients retained a strong marginal FC backbone (Fig. 2a, b).

**Figure 2.**
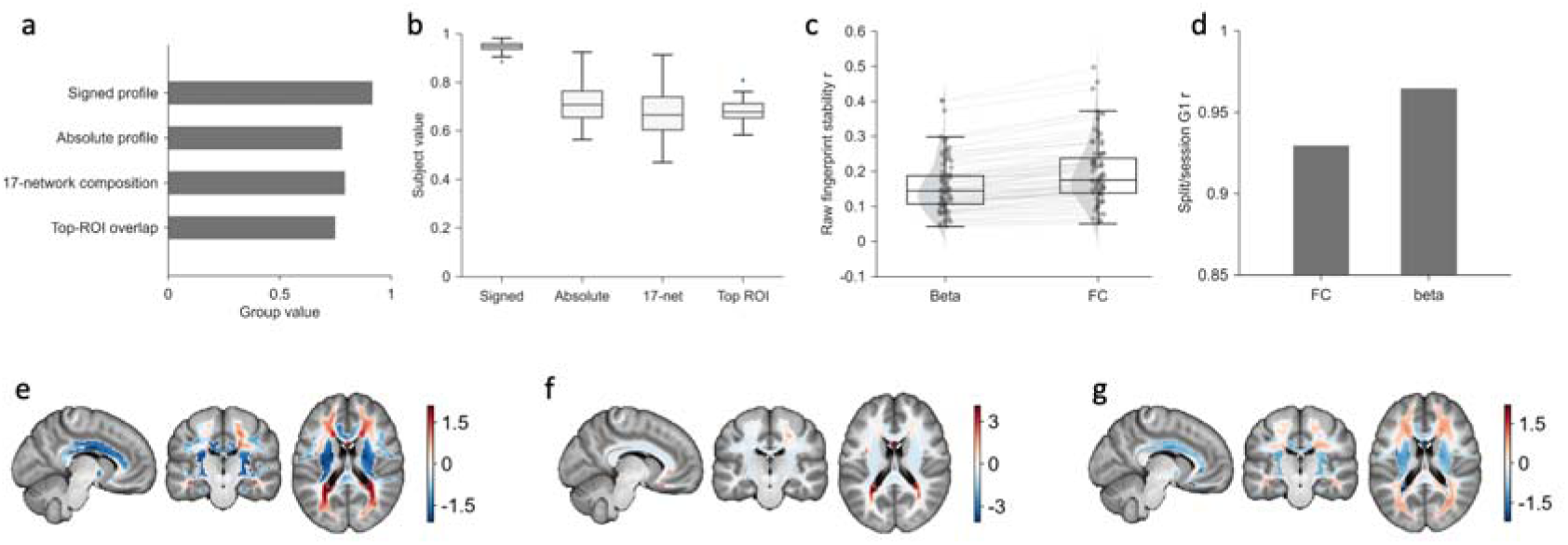
Ridge beta fingerprints recapitulate but diverge from marginal GM-WM functional connectivity. (a) Group-level correspondence between ridge beta fingerprints and marginal FC fingerprints, summarized by signed profile similarity, absolute profile similarity, 17-network composition similarity, and top-ROI overlap. (b) Subject-level distributions of the same beta-FC correspondence metrics, showing consistent similarity across individuals. (c) Split/session stability of raw beta and FC fingerprints. Points and boxes show subject-level stability estimates, and grey lines indicate paired values from the same subject. (d) Split/session stability of the first gradient (G1) derived from beta and FC fingerprints. (e) Fixed beta G1 template, summarizing the dominant spatial axis of ridge-derived cortical encoding profiles across WM voxels. (f) Fixed FC G1 template after sign alignment to the beta G1 template. Because the gradient sign is arbitrary, the FC template was sign-aligned before visualization. (g) Spatial difference between beta G1 and sign-aligned FC G1. Warm colors indicate a relatively higher beta-G1 position, whereas cool colors indicate a relatively higher FC-G1 position.

Despite this shared backbone, beta fingerprints and FC fingerprints differed in their reliability structure and low-dimensional organization. Raw high-dimensional FC fingerprints were more stable than raw beta fingerprints across session splits (mean r = 0.198 for FC versus 0.158 for beta), consistent with the additional variance introduced by conditional multivariate estimation. However, the first beta gradient was highly reproducible across splits (G1 r = 0.965) and slightly more stable than the first FC gradient (G1 r = 0.929). Fixed beta and FC G1 templates were related but not identical (template r = 0.701), and their spatial difference defined a beta-minus-FC G1 axis that captured how conditional encoding profiles diverged from marginal FC profiles (Fig. 2c-g).

This low-dimensional axis was treated as a continuous organizational gradient rather than a discrete WM parcellation. K-means clustering of beta fingerprints showed weak silhouette support for both k = 7 and k = 17 (mean silhouette = 0.219 and 0.174, respectively), arguing against a strong discrete clustering interpretation and motivating the gradient-based analyses used below (Fig. S5a).

### Local prediction residuals reveal organization beyond marginal FC strength

Because ridge beta fingerprints were related to marginal FC but not reducible to it, we tested whether local prediction accuracy also contained structure beyond local FC magnitude. Local z-scored R^2^ and z-scored FC RMS maps showed related but visibly distinct spatial patterns. After regressing local zR^2^ on local zFC RMS within each subject, the resulting R^2^ residual map highlighted WM regions where prediction was higher or lower than expected from marginal FC strength alone (Fig. 3a-c).

**Figure 3.**
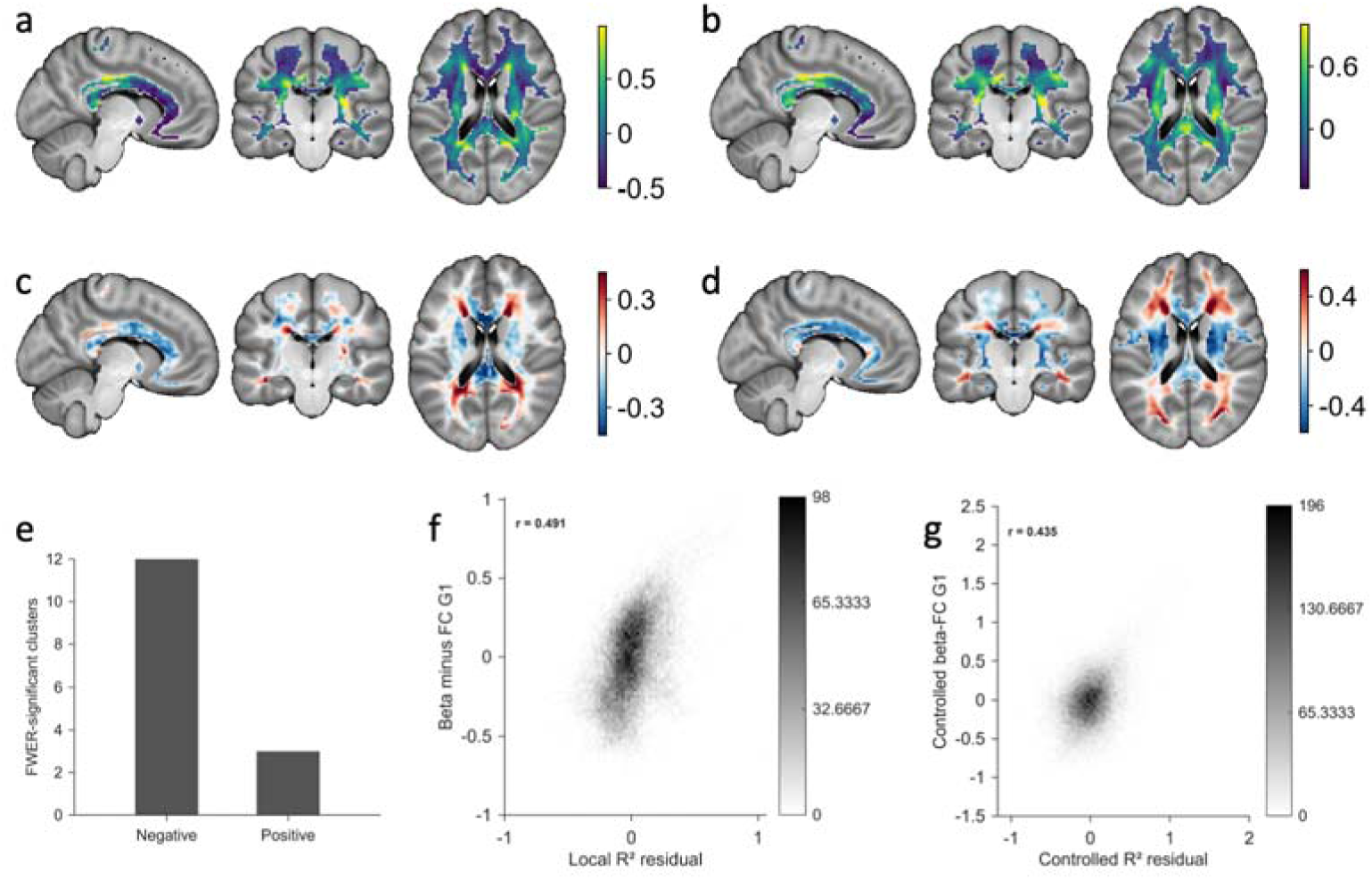
Local prediction residuals reveal a beta-FC divergence axis beyond marginal functional connectivity. (a) Local z-scored cross-validated R² map, showing the spatial distribution of cortical-to-WM prediction accuracy. (b) Local z-scored FC RMS map, summarizing the magnitude of marginal GM-WM functional connectivity for each WM voxel. (c) R² residual after accounting for local FC magnitude, highlighting WM regions where encoding-based predictability exceeds or falls below that expected from marginal FC strength. (d) Subject-mean beta-FC G1 difference map, reflecting the spatial divergence between the dominant ridge beta gradient and the corresponding FC gradient. (e) FWER-significant signed residual clusters, summarized separately for negative and positive residual effects. (f) Voxel-wise density plot showing the association between local R² residuals and beta-FC G1 differences before nuisance control. The color scale indicates voxel count per bin. (g) Voxel-wise density plot showing the association between controlled R² residuals and controlled beta-FC G1 differences after nuisance adjustment. The color scale indicates voxel count per bin.

The local residual map showed statistically reliable signed spatial structure. Cluster-mass permutation identified 15 FWER-significant signed clusters, including 3 positive clusters comprising 3,186 voxels and 12 negative clusters comprising 1,332 voxels. The largest positive clusters contained 1,961 and 1,203 voxels, whereas the largest negative cluster contained 880 voxels. These residual effects indicate that the cortical encoding model captured spatially organized WM predictability not fully summarized by local FC magnitude (Fig. 3e).

The FC-adjusted prediction residuals aligned with the beta-minus-FC G1 axis. The spatial correlation between the group R^2^ residual map and the subject-mean beta-minus-FC G1 map was r = 0.491, indicating that WM voxels with stronger conditional beta-FC divergence tended to show higher-than-expected prediction after accounting for marginal FC strength (Fig. 3d,f). This convergence linked the local accuracy residual and the fingerprint-gradient analyses as two views of the same conditional encoding axis.

### Spatial and signal-quality controls support a WM-specific effect

We then evaluated whether the residual-G1 relationship could be explained by simple spatial or signal-quality covariates. Nuisance maps included distance to cortical GM, distance to the WM mask boundary, WM prevalence across subjects, temporal signal standard deviation, and spatial coordinate terms (Fig. 4a-d). After residualizing both the R^2^ residual map and beta-minus-FC G1 map against this nuisance design, their spatial correlation remained substantial (r = 0.435), only modestly reduced from the uncontrolled value of r = 0.491 (Fig. 4e).

**Figure 4.**
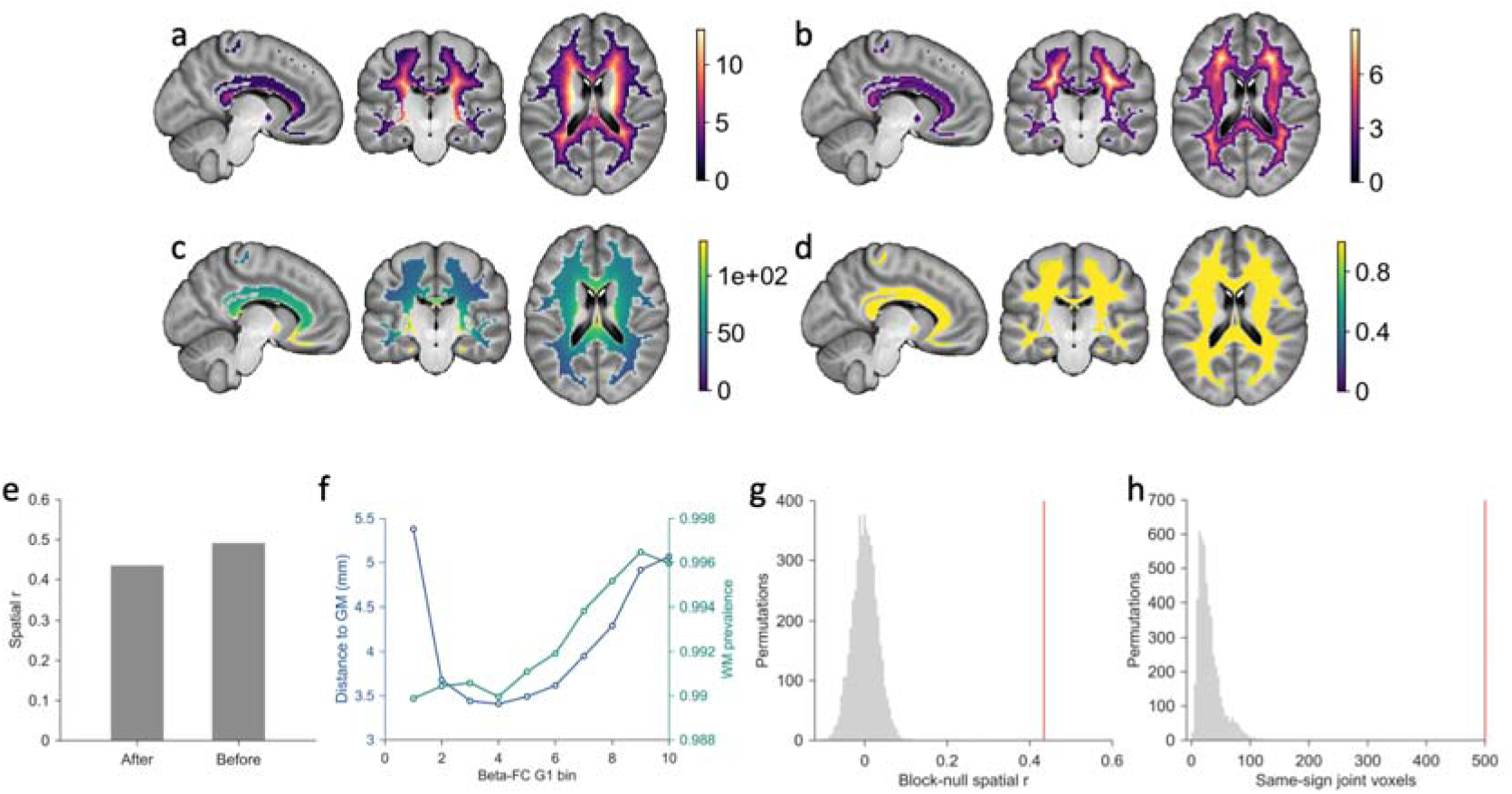
Spatial and signal-quality controls for the beta-FC divergence axis. (a) Distance from each WM voxel to cortical GM, used to assess potential GM contamination. (b) Distance from each WM voxel to the WM mask boundary, used to characterize proximity to mask edges and partial-volume-prone regions. (c) Temporal standard deviation map, summarizing local BOLD signal variability across WM voxels. (d) WM prevalence map, showing the proportion of subjects in which each voxel was classified as WM. (e) Effect of nuisance control on the spatial correspondence between the R² residual map and the beta-FC G1 difference map. Bars show the spatial correlation before and after adjustment for spatial and signal-quality covariates. (f) G1-bin contamination diagnostics. Mean distance to cortical GM and WM prevalence are plotted across beta-FC G1 bins to evaluate whether the divergence axis reflects systematic variation in GM proximity or WM mask prevalence. (g) Spatial-block null distribution for the controlled map correlation. The red vertical line indicates the observed spatial correlation relative to the block-permuted null distribution. (h) Spatial-block null distribution for same-sign joint voxels. The red vertical line indicates the observed number of joint voxels relative to the block-permuted null distribution.

Voxelwise controlled inference showed that both components of this relationship remained spatially extensive after nuisance adjustment. The controlled R^2^ residual map contained 994 FWER-significant voxels, the controlled beta-minus-FC G1 map contained 2,731 FWER-significant voxels, and the controlled zR^2^ map after adjustment for zFC and spatial confounds contained 1,026 FWER-significant voxels. Their same-sign joint effect included 513 voxels, with 97.5% sign agreement between the two controlled effects (Fig. S8).

The result also survived a spatial-autocorrelation-preserving volumetric block null. Using 12-mm spatial blocks, 553 WM blocks, and 5,000 permutations, the controlled residual-G1 map correlation exceeded the block-permuted null (block p = 0.0002). The controlled joint voxel count and same-sign joint voxel count also exceeded the block null (both p = 0.0002), indicating that the effect was not an artifact of smooth spatial autocorrelation alone (Fig. 4g,h). Finally, G1-bin contamination diagnostics showed that the high-G1/high-residual end was not simply closer to cortical GM; the highest G1 bin had mean distance to GM of 5.06 mm and WM prevalence of 0.996, whereas the lowest G1 bin had mean distance to GM of 5.38 mm and WM prevalence of 0.990 (Fig. 4f).

### The beta-FC divergence axis organizes networks, tracts, and validation results

To interpret the conditional encoding axis, we ordered WM voxels into ten equal-count bins along beta-minus-FC G1. Across bins, marginal FC strength and FC-adjusted prediction residuals dissociated. The low-G1 end showed relatively high FC magnitude but negative residual prediction (bin 1: mean zFC RMS = 0.525; mean R^2^ residual = -0.094), whereas the high-G1 end showed low FC magnitude but positive residual prediction (bin 10: mean zFC RMS = -0.148; mean R^2^ residual = 0.239). This pattern persisted after spatial and signal-quality controls, with bin 10 retaining a controlled residual of 0.239 and zR^2^ after zFC plus confound adjustment of 0.242 (Fig. 5a-c).

**Figure 5.**
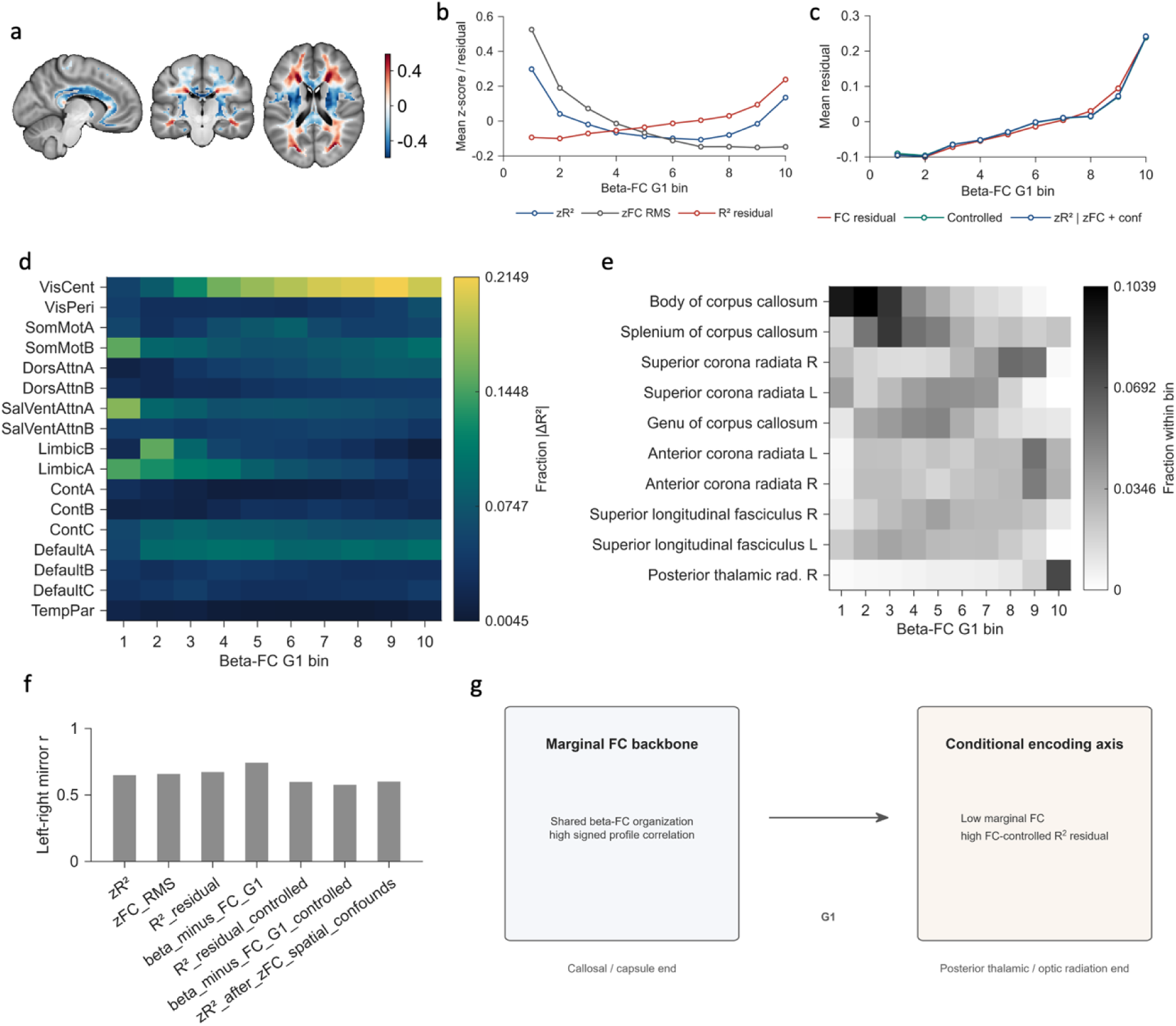
The beta-FC divergence axis organizes prediction residuals, cortical network contributions and WM tract anatomy. (a) Beta-FC G1 axis map used to order WM voxels into ten bins. Warm and cool colors indicate opposite ends of the divergence axis between ridge beta gradients and marginal FC gradients. (b) Prediction and connectivity structure across beta-FC G1 bins. Lines show mean z-scored R², z-scored FC RMS and R² residual values within each bin, illustrating how encoding accuracy, marginal FC magnitude and FC-adjusted prediction residuals vary along the axis. (c) Controlled residual profiles across beta-FC G1 bins. Raw FC residuals, nuisance-controlled residuals and zR² after adjustment for FC and spatial confounds are plotted to show that the residual organization persists after covariate control. (d) Cortical network contribution across beta-FC G1 bins. The heatmap shows the fractional contribution of each Yeo 17-network predictor group to the absolute change in prediction accuracy. (e) JHU tract composition across beta-FC G1 bins. The heatmap shows the fraction of voxels within each bin assigned to major WM tracts, linking the functional divergence axis to tract anatomy. (f) Bilateral organization of major maps. Bars show left-right mirror correlations for prediction, FC, residual and controlled maps, indicating the degree of hemispheric symmetry. (g) Conceptual model. The dominant GM-WM organization reflects a marginal FC backbone, whereas the beta-FC G1 axis reveals a conditional encoding dimension characterized by relatively low marginal FC but high FC-controlled prediction residuals toward the posterior thalamic/optic-radiation end.

Cortical-network ablation profiles changed systematically along the same axis. Low-G1 bins were dominated by salience/ventral-attention, somatomotor, and limbic contributions, whereas high-G1 bins showed larger visual-central and default-network contributions. In bin 10, visual-central predictors accounted for the largest fraction of absolute prediction change (0.198), followed by default A (0.101) and somatomotor B (0.099), linking the high residual end of the axis to a distinct cortical contribution profile (Fig. 5d).

The axis also mapped onto tract anatomy. Low-G1 bins contained larger fractions of corpus callosum, internal capsule, and external capsule voxels, whereas the high-G1 end was enriched for posterior thalamic radiation including optic radiation, posterior corona radiata, and sagittal stratum. Direct JHU tract summaries supported the same anatomical organization: tract-level mean R^2^ residuals correlated with tract-level mean beta-minus-FC G1 values (r = 0.658). The most positive residual tracts included tapetum, posterior thalamic radiation, posterior corona radiata, and sagittal stratum, while the most negative tracts included body and genu of corpus callosum and internal-capsule segments (Fig. 5e; Fig. S6).

Several additional analyses supported reproducibility while clarifying the limits of interpretation. Major maps showed bilateral mirror organization, with left-right correlations ranging from 0.649 for zR^2^ to 0.743 for beta-minus-FC G1; controlled maps also retained moderate symmetry (Fig. 5f). Cross-session validation showed strong group-level reproducibility for both the local R^2^ residual map (REST1-REST2 group map r = 0.741) and the beta-minus-FC G1 map (cross-session group map r = 0.955), although subject-level cross-session correlations were lower, as expected for noisier individual WM maps (Fig. S7). Exploratory behavior-prediction analyses did not reveal robust corrected associations: global encoding features showed no significant corrected incremental prediction after confounds, and all local map-by-outcome models had corrected q_delta_R^2^ = 1.0 (Fig. S5b,c). Together, these analyses support a model in which GM-WM organization contains a broad marginal FC backbone plus a conditional encoding axis that is most evident in posterior thalamic/optic-radiation and related posterior WM territories (Fig. 5g).

## DISCUSSION

In this study, we used a cortical encoding framework to characterize how distributed GM activity predicts spontaneous BOLD fluctuations in strict cerebral WM. Three main findings emerged. First, cortical parcel activity predicted held-out WM BOLD signals across a broad portion of the WM mask, with modest but reliable cross-validated accuracy. This effect was spatially organized, was stronger for fine-grained cortical parcels than for coarse network averages, and exceeded circular-shift null expectations, indicating that it was not simply a consequence of temporal autocorrelation. Second, ridge-derived beta fingerprints strongly recapitulated conventional marginal gray-to-white functional connectivity fingerprints, demonstrating that conditional encoding and marginal FC share a common large-scale functional backbone. Third, the divergence between the ridge beta gradient and the FC gradient revealed a reproducible organizational axis along which local prediction accuracy dissociated from marginal FC magnitude. This beta-FC divergence axis organized FC-adjusted prediction residuals, cortical network contribution profiles, and WM tract anatomy, suggesting that multivariate cortical encoding captures a dimension of WM functional organization that is not fully described by pairwise GM-WM correlations.

A central implication of these findings is that marginal functional connectivity provides an important but incomplete description of gray-to-white functional coupling. Traditional GM-WM FC asks whether a given cortical parcel and a given WM voxel fluctuate together. This pairwise view is valuable, but it is intrinsically sensitive to broad cortical synchrony and collinearity among large-scale networks. In contrast, the ridge encoding model asks a different question: how well can the distributed cortical activity pattern predict a local WM signal after accounting for shared variance among cortical predictors? The answer was not independent of marginal FC; indeed, beta and FC fingerprints were highly similar. However, the residual difference between them was spatially structured and reproducible. Thus, the present results support a complementary framework in which marginal FC captures the dominant synchronization backbone of GM-WM coupling, whereas conditional encoding reveals how distributed cortical systems jointly account for local WM dynamics.

The strong similarity between ridge beta fingerprints and FC fingerprints is itself informative. It indicates that the multivariate model does not generate an arbitrary or method-specific representation of WM organization. Rather, the dominant encoding structure remains anchored to the same large-scale GM-WM coupling patterns observed with conventional FC. At the same time, the first beta gradient and the first FC gradient were related but not identical, and their difference defined a stable beta-FC divergence axis. This pattern suggests that the most interpretable information in the encoding model may not lie in raw high-dimensional beta coefficients, which are sensitive to regularization and predictor collinearity, but in low-dimensional, reproducible axes of coefficient organization. In this sense, the beta-FC G1 axis provides a compact representation of where conditional cortical encoding agrees with, or diverges from, marginal functional connectivity.

The local prediction residual analyses provide converging evidence for this interpretation. If cortical-to-WM prediction were merely a restatement of local FC strength, then prediction accuracy should be fully explained by the magnitude of marginal GM-WM correlations. Instead, after regressing local prediction accuracy on local FC magnitude, the residual map showed reliable spatial organization. Moreover, these FC-adjusted prediction residuals aligned with the beta-FC G1 difference map. WM regions at one end of the axis showed relatively strong marginal FC but lower-than-expected prediction residuals, whereas regions at the opposite end showed relatively weak marginal FC but higher-than-expected prediction residuals. This dissociation suggests that some WM signals may not be dominated by strong pairwise synchronization with cortex, but may nevertheless be predictable from distributed cortical activity patterns when considered jointly.

Anatomically, the beta-FC divergence axis separated callosal and capsular regions from posterior thalamic, optic-radiation, posterior corona-radiata, and sagittal-stratum regions. The low-G1 end was enriched for corpus callosum and internal/external capsule voxels and was characterized by relatively high marginal FC but negative FC-adjusted prediction residuals. This pattern may reflect WM regions whose BOLD fluctuations are strongly synchronized with broad cortical activity but do not gain as much additional explanatory structure from conditional multivariate modeling. In contrast, the high-G1 end was enriched for posterior thalamic radiation, optic radiation, posterior corona radiata, and sagittal stratum, and showed relatively low marginal FC but positive FC-adjusted prediction residuals. This suggests that posterior projection and association pathways may occupy a functional regime in which distributed cortical activity predicts local WM dynamics better than expected from marginal FC strength alone. Importantly, this interpretation does not require that these tracts have stronger overall coupling to cortex; rather, it suggests that their coupling may be more conditionally distributed across cortical systems.

The cortical network ablation results further support the view that the divergence axis reflects changes in cortical contribution profiles rather than a purely local WM property. Along the beta- FC G1 axis, the relative contribution of cortical networks changed systematically, with lower-G1 bins showing stronger salience/ventral-attention, somatomotor, and limbic contributions, and higher-G1 bins showing greater visual-central and default-network contributions. These results should not be interpreted as causal evidence that specific cortical networks drive specific WM tracts, because cortical networks are mutually correlated and ridge coefficients distribute shared variance across predictors. Nevertheless, the systematic organization of ablation profiles indicates that the beta-FC divergence axis is linked to differences in distributed cortical predictive structure. This provides a bridge between voxelwise WM BOLD organization, large-scale cortical networks, and tract anatomy.

A major methodological concern in WM fMRI is that apparent WM effects may reflect GM contamination, partial-volume effects, vascular artifacts, motion, or spatial smoothing. Several aspects of the present design reduce these concerns. The analysis used a strict WM95 mask with no overlap with the cortical atlas or group GM mask, and WM smoothing was constrained within the WM mask. Physiological and motion covariates were regressed from the time series before modeling. The prediction effect survived circular-shift null testing, indicating that it was not explained by autocorrelated time series alone. The residual-G1 relationship also persisted after controlling for distance to cortical GM, distance to the WM boundary, WM prevalence, temporal standard deviation, and spatial coordinate terms. Finally, the controlled map correlation and joint voxel effects exceeded a spatial-block permutation null, arguing against smooth spatial autocorrelation as a sufficient explanation. These controls do not prove that the observed signals are purely neural, but they support the conclusion that the conditional encoding axis is a reproducible WM-specific organization not reducible to simple anatomical proximity or signal-quality gradients.

The present work also has implications for how WM BOLD should be analyzed. Much previous work has used FC, clustering, or network-level summaries to describe intrinsic WM functional organization. Our results suggest that encoding models can complement these approaches by separating shared marginal synchronization from conditional predictive structure. This is especially relevant for WM because local BOLD signals may reflect many-to-one relationships with distributed cortical systems. In this context, pairwise FC may identify regions that are strongly synchronized with cortex, whereas ridge encoding may identify regions whose fluctuations are best explained by a distributed cortical state. This distinction may be useful for future studies of development, aging, disease, or task engagement, where WM abnormalities may manifest not only as changes in FC magnitude but also as changes in the conditional mapping between cortical activity and WM dynamics.

Several limitations should be acknowledged. First, although prediction accuracy was reliable and spatially structured, the absolute cross-validated R² values were modest. This is expected given the lower signal-to-noise ratio of WM BOLD and the conservative held-out prediction framework, but it cautions against overstating the magnitude of explained variance. Second, ridge beta coefficients are not direct measures of causal influence or effective connectivity. Because cortical predictors are correlated, beta weights and ablation scores should be interpreted as conditional predictive summaries rather than independent biological contributions. Third, fMRI remains an indirect measure of neural and vascular activity, and the present data cannot fully distinguish neural, vascular, metabolic, and residual physiological sources of WM BOLD fluctuations. Fourth, the study focused on young healthy adults from the HCP dataset; independent replication, lifespan samples, clinical cohorts, and task-based datasets will be needed to determine whether the beta-FC divergence axis generalizes across populations and brain states. Fifth, although the lagged model produced only a small improvement over the zero-lag model, the present analysis was not designed to fully characterize regional hemodynamic delay or directionality. Future work combining encoding models with explicit hemodynamic-lag estimation may clarify how timing differences contribute to conditional GM-WM coupling. Finally, tract interpretation was based on atlas overlap and should be refined using subject-specific diffusion anatomy and higher-resolution acquisitions.

In conclusion, this study shows that spontaneous WM BOLD activity can be modestly but reliably predicted from distributed cortical GM activity. Ridge beta fingerprints largely preserve the marginal GM-WM FC backbone, but their divergence from FC reveals a stable conditional encoding axis that organizes prediction residuals, cortical network contributions, and tract anatomy. These findings support a view of WM as an active component of large-scale functional organization and demonstrate that multivariate cortical encoding can reveal aspects of GM-WM coupling that are not captured by conventional pairwise functional connectivity alone.

## Data Availability

The HCP-Y data used in this study are available in the HCP database https://www.humanconnectome.org/

## Code Availability

The custom scripts and the computational algorithms developed for the neuroimaging analysis in this study are freely available in the public GitHub repository at: https://github.com/geyerou/Engagement-by-regression

Detailed instructions for reproducing the key results are provided in the repository’s README file.

Other software and toolboxes that are required for running the code include: Nilearn: https://nilearn.github.io/

SPM12: https://www.fil.ion.ucl.ac.uk/spm/software/spm12/

TAPAS PhysIO: https://github.com/ComputationalPsychiatry/PhysIO

BrainSpace: https://github.com/MICA-MNI/BrainSpace

## ACKNOWLEDGMENTS

This work was supported by the National Institutes of Health (NIH) grants R01 NS113832 (JG) and R01 NS129855 (ZD).

Data of young adults were provided by the Human Connectome Project, WU-Minn Consortium (Principal Investigators: David Van Essen and Kamil Ugurbil; 1U54MH091657), funded by the 16 NIH Institutes and Centers that support the NIH Blueprint for Neuroscience Research; and by the McDonnell Center for Systems Neuroscience at Washington University.

## AUTHOR CONTRIBUTIONS

M.L.: Writing, Visualization, Validation, Software, Methodology, Investigation, Conceptualization. Z.D.: Supervision and Funding acquisition. J.C.G: Supervision and Funding acquisition.

