## Supplemental Figures for "White-Matter BOLD Encoding Beyond Marginal Connectivity"

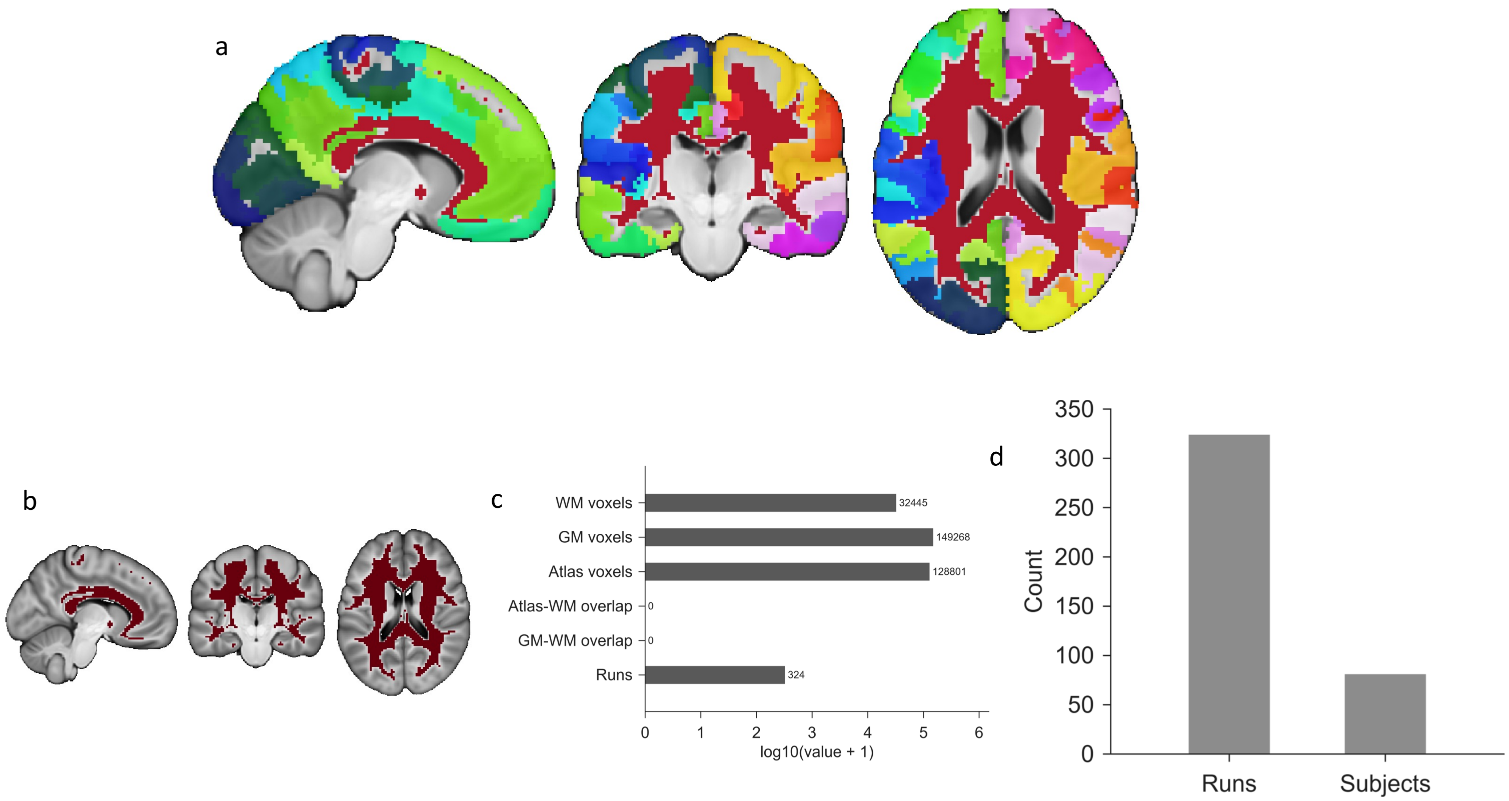


Figure S1. Anatomical masks, gray-white matter separation, and included data. (a) Combined anatomical context showing the Schaefer-400 cortical GM atlas together with the strict WM95 target mask. Cortical parcels are displayed in distinct atlas colors, and the WM95 mask is shown in red. (b) Strict WM95 mask alone, illustrating the WM voxel space used as the prediction target. (c) Mask and run quality-control summary. Counts are shown on a log10 (value + 1) scale; the zero atlas-WM and GM-WM overlap values confirm separation between cortical GM predictors and WM target voxels. (d) Included data, showing the number of resting-state runs and subjects retained for analysis.

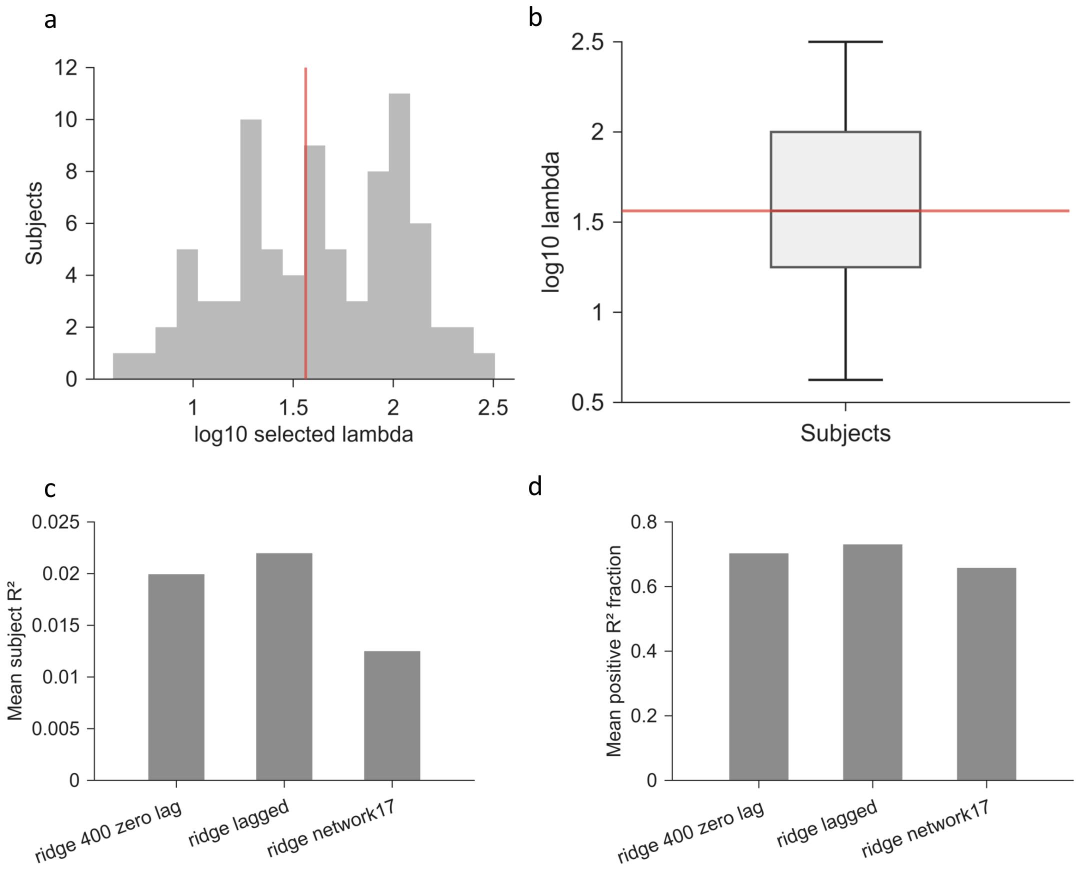


Figure S2. Ridge-regularization parameters and model-sensitivity summaries. (a) Distribution of subject-specific ridge-regularization parameters selected during cross-validation. The red vertical line indicates the shared lambda reference used for downstream group-level analyses. (b) Boxplot of subject-specific selected lambda values, shown on a log10 scale. The red horizontal line indicates the shared lambda reference. (c) Mean subject-level prediction accuracy across model variants, including the zero-lag Schaefer-400 ridge model, the lagged ridge model, and the reduced 17-network ridge model. (d) Mean fraction of WM voxels with positive cross-validated $R^2$ across the same model variants, summarizing the spatial extent of above-zero prediction performance.

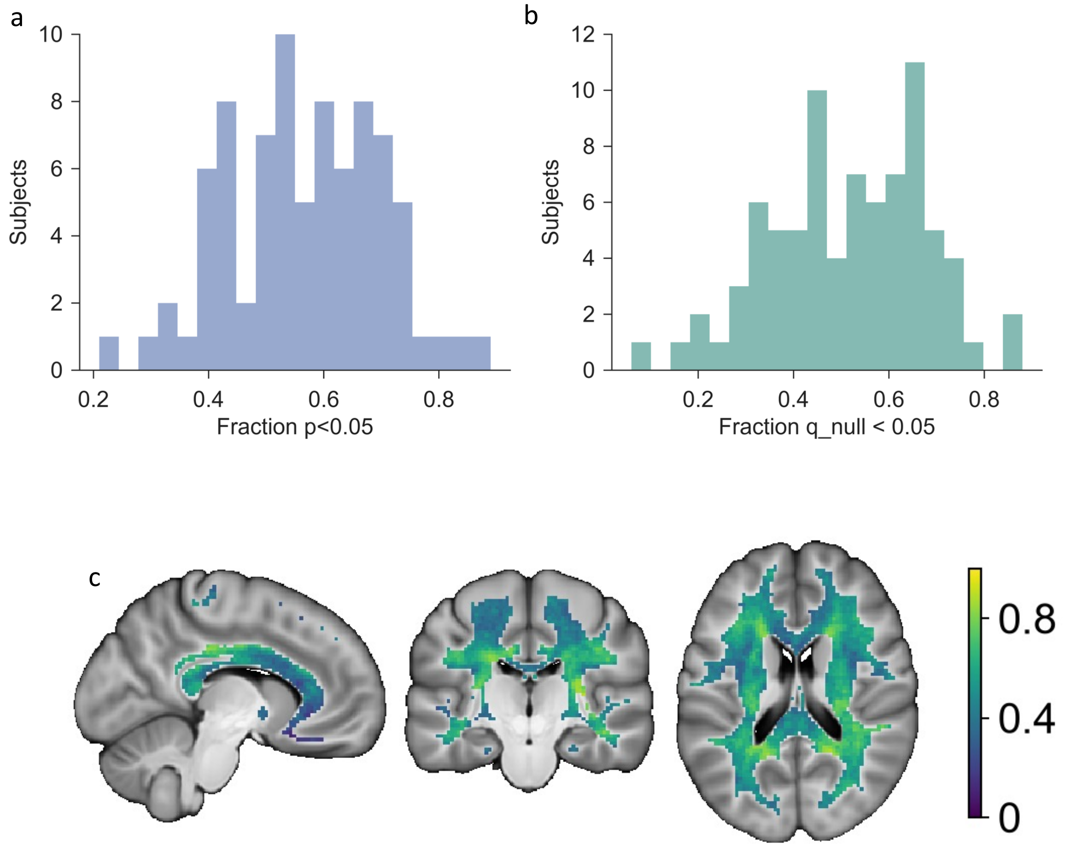


Figure S3. Subject-level and spatial extent of above-null WM prediction. (a) Distribution across subjects of the fraction of WM voxels whose observed prediction accuracy exceeded the circular-shift null distribution at the voxel-wise uncorrected threshold. (b) Distribution across subjects of the fraction of WM voxels surviving FDR correction against the circular-shift null distribution (q_null < 0.05). (c) Spatial consistency of FDR-corrected above-null prediction across subjects. The map shows, for each WM voxel, the fraction of subjects in which the voxel survived the q_null < 0.05 threshold; the color scale ranges from 0 to 1.

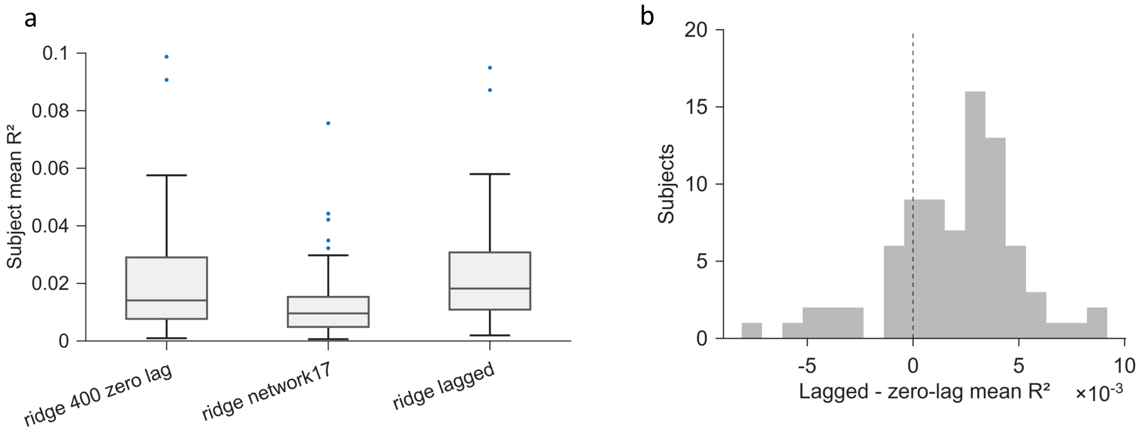


Figure S4. Model sensitivity and lagged-model comparison. (a) Subject-level mean prediction accuracy across model variants, including the zero-lag Schaefer-400 ridge model, the reduced 17-network ridge model, and the lagged ridge model. Boxplots show the distribution of mean WM $R^2$ values across subjects, with points indicating individual subjects. (b) Distribution of the within-subject difference in mean prediction accuracy between the lagged and zero-lag ridge models. The dashed vertical line marks zero difference; positive values indicate higher prediction accuracy for the lagged model.

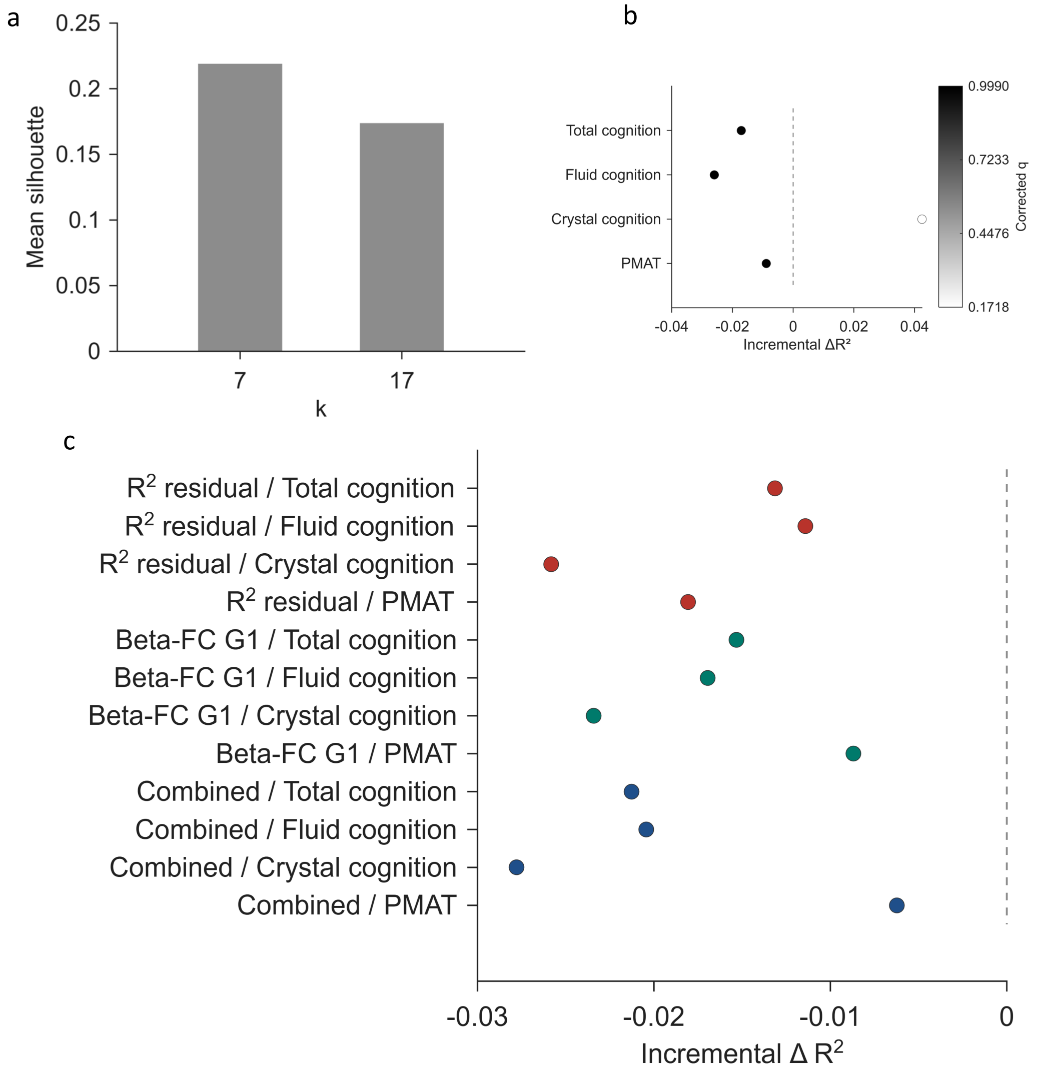


Figure S5. Clustering support and behavioral-prediction sensitivity analyses. (a) K-means silhouette analysis comparing two candidate clustering solutions. Bars show the mean silhouette value for k = 7 and k = 17. (b) Global behavioral-prediction analysis. Points show the incremental cross-validated $\Delta R^2$ for predicting cognitive outcomes after adding global WM encoding features to the baseline model. The dashed vertical line marks zero incremental prediction, and point shading indicates the corrected q value. (c) Local behavioral-prediction analysis across map-by-outcome combinations. Incremental $\Delta R^2$ values are shown for models using the $R^2$ residual map, the beta-FC G1 map, or their combined feature set to predict total cognition, fluid cognition, crystal cognition, and PMAT performance. The dashed vertical line marks zero incremental prediction.

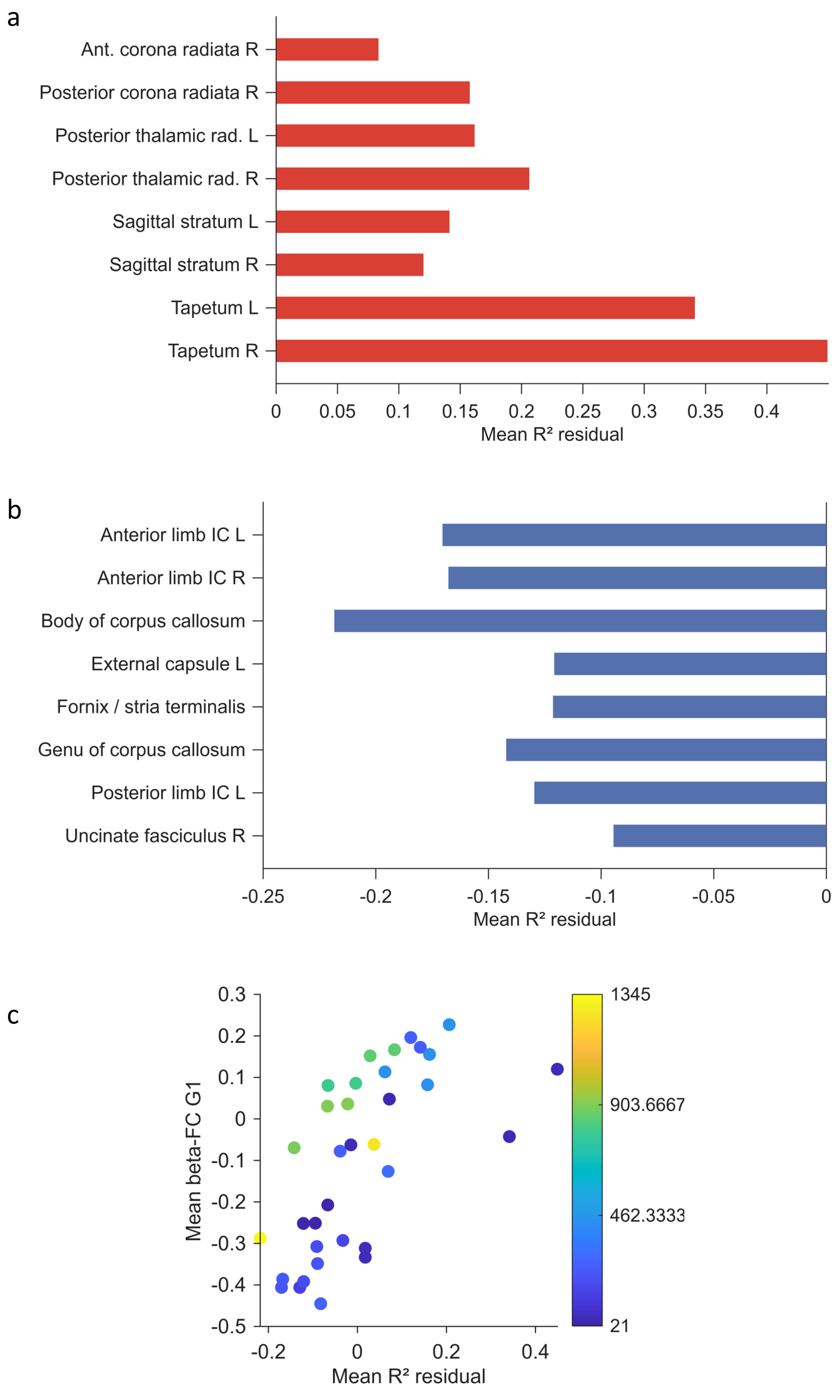


Figure S6. Tract-level organization of FC-adjusted prediction residuals and beta-FC divergence. (a) JHU WM tracts with the largest positive mean $R^2$ residuals after accounting for local FC magnitude. Positive values indicate tracts in which cortical-to-WM prediction accuracy was higher than expected from marginal GM-WM FC strength. (b) JHU WM tracts with the largest negative mean $R^2$ residuals. Negative values indicate tracts in which prediction accuracy was lower than expected from marginal GM-WM FC strength. (c) Tract-level convergence between mean $R^2$ residuals and mean beta-FC G1 values. Each point represents a JHU tract, and the point color indicates the number of voxels assigned to that tract.

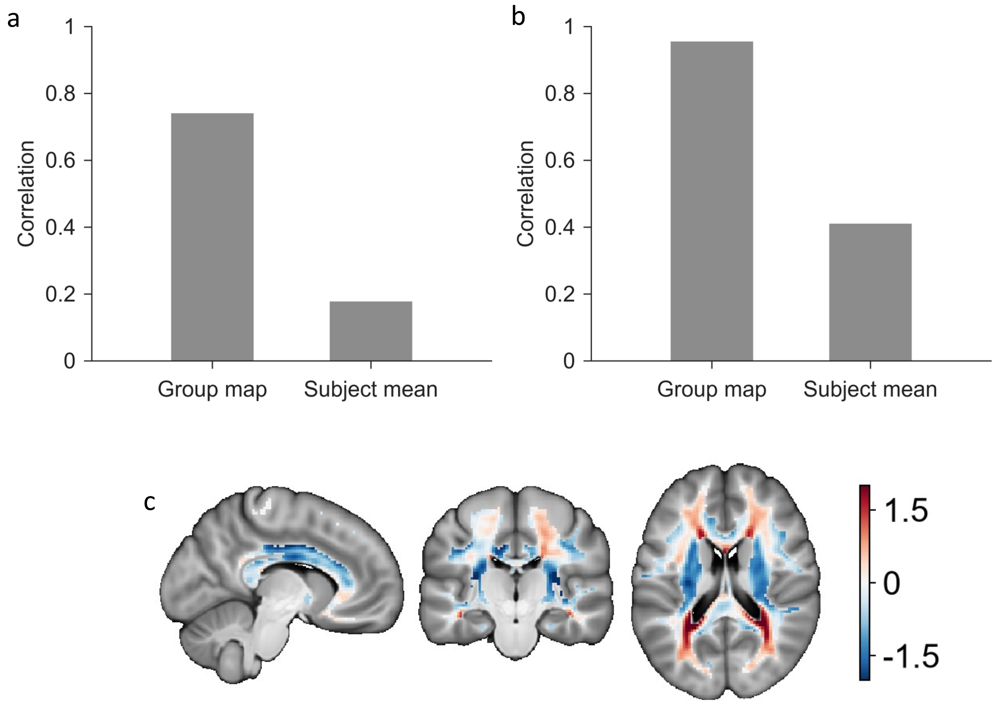


Figure S7. Cross-session validation of prediction residuals and the beta-FC divergence axis. (a) REST1-to-REST2 validation of the $R^2$ residual map. Bars show the spatial correlation between session-specific group maps and the mean subject-level cross-session correlation. (b) REST1-to-REST2 validation of the beta-FC G1 map. Bars show the spatial correlation between session-specific group maps and the mean subject-level cross-session correlation. (c) Cross-session beta-FC G1 map, showing the reproducible spatial organization of the beta-FC divergence axis across independent resting-state sessions.

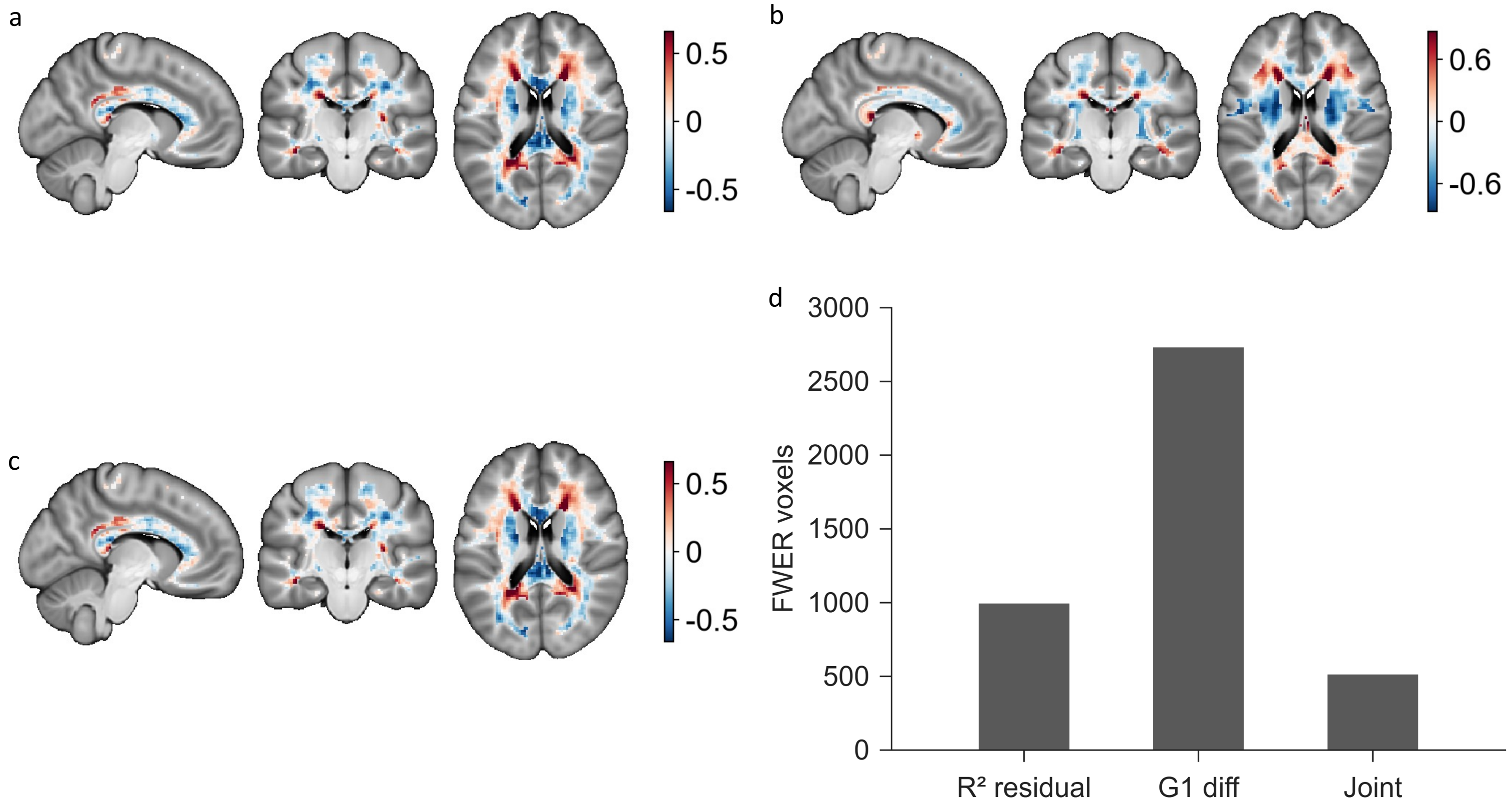


Figure S8. Controlled spatial maps and voxel-wise extent of residual beta-FC effects. (a) Controlled $R^2$ residual map after adjustment for local FC magnitude and nuisance covariates, showing WM regions where cortical-to-WM prediction accuracy remained higher or lower than expected after control. (b) Controlled beta-FC G1 difference map after nuisance adjustment, showing the residual spatial divergence between ridge beta gradients and marginal FC gradients. (c) Controlled $zR^2$ map after adjustment for zFC and spatial confounds, showing prediction-related variation that remains after accounting for connectivity magnitude and anatomical covariates. (d) FWER-corrected voxel counts for the controlled $R^2$ residual, controlled G1 difference, and their same-sign joint effect.
